# sc4D: spatio-temporal single-cell transcriptomics analysis through embedded optimal transport identifies joint glial response to Alzheimer’s disease pathology

**DOI:** 10.1101/2025.11.19.689166

**Authors:** Ishir Rao, Manolis Kellis, Yosuke Tanigawa

## Abstract

A precise understanding of disease-associated spatio-temporal transcriptional dynamics is critical for nominating therapeutic interventions that drive desired perturbation responses. In disease contexts, pathological processes unfold across diverse cellular states, spatial environments, and timescales; however, current computational approaches have a limited ability to jointly model these complex dynamics or infer cellular trajectories from *in silico* perturbation experiments. Here, we present sc4D, a biologically interpretable spatio-temporal (4D) analysis framework for single-cell transcriptomics in disease studies, integrating autoencoder embeddings with optimal transport. Applying sc4D to longitudinal spatial transcriptomics samples from Alzheimer’s disease mouse models, we report known disease biology and novel, testable mechanisms, including late-stage microglia-astrocyte syncytium near amyloid-β plaques. *In silico* perturbations predict the restoration of protective, anti-inflammatory microglia through CAMTA1 activation or Donepezil, as validated by independent experimental findings. Overall, our work highlights the critical benefits of joint spatio-temporal modeling for elucidating disease mechanisms and predicting candidate interventions to improve cellular response.

## Introduction

Alzheimer’s disease (AD) is a progressive neurodegenerative disease and a leading cause of dementia in the elderly. AD is often characterized by several pathological aggregates, including amyloid beta (Aβ) plaque and paired helical filament (PHF+) tau tangles[1–3], that disrupt neuronal function and ultimately lead to cell death. AD has been documented to display significant spatial heterogeneity[4–7], specifically in spatially enriched subpopulations of glial cell types[8,9] and differentially expressed genes[10–12] surrounding plaque. Moreover, the gradual progression of AD over seven to ten years on average[13] necessitates longitudinal analysis to capture all stages of disease response.

To study the spatial and longitudinal landscape of AD, several pioneering studies have started to investigate the molecular changes in mouse AD models that display accelerated AD features. One such example is the TauPS2APP mode, which, at 8 months of age, displays Aβ plaque and tau deposition and, at 13 months of age, presents an advanced stage of disease with severe neurodegeneration and neuroinflammation[14]. In particular, spatially-resolved single-cell RNA sequencing (scRNA-seq) offers unprecedented insight into the spatial heterogeneity of gene expression across cell types in disease contexts[15,16]. Despite the advancements in experimental characterization, critical computational challenges remain to be solved to understand the cellular response to AD by capturing both spatial organization and longitudinal trends of cells in complex tissues.

Several recent advances have begun investigating this spatio-temporal landscape and have yielded new insights into cellular response to localized stressors. For spinal cord injury[17], for example, spatio-temporal analysis revealed distinct microglia and astrocyte subtypes that appear at later stages of response. Similarly, for colorectal cancer[18], spatio-temporal analysis revealed that CD8+ T-cells alter their transcriptome surrounding tumors over time in response to drug perturbations. Due to the destructive nature of transcriptome profiling on the cell, however, both studies required spatial transcriptomics measurements at a high temporal resolution, which may not be feasible for biological processes at larger timescales. As a result, techniques are necessary to predict longitudinal trends from sparse time-series single-cell data.

One such approach, trajectory inference, models cellular and transcriptional dynamics over time and has been most extensively applied in developmental contexts[19]. Methods such as Monocle3[20] and PAGA[21] apply low-dimensional projections through t-distributed Stochastic Neighbor Embedding (t-SNE) or Principal Component Analysis (PCA) as preprocessing. Then, computational techniques like diffusion maps[22] or continuous vector fields[23] are implemented to predict trajectories. Optimal transport (OT), a mathematical framework to find the most efficient mapping between two distributions, is another method for trajectory inference that integrates time-series single-cell data[24]. Techniques such as Waddington-OT[25] and TIGON[23] have applied optimal transport in the context of Waddington’s epigenetic landscape[26,27] and single-cell developmental trajectories. More recently, methods such as Moscot[28,29] and DeST-OT[28] have implemented OT for spatio-temporal alignment of scRNA-seq data in developmental contexts. In addition, techniques like CellOT[30] have implemented OT to predict transcriptome response to perturbations. Specifically in disease contexts, ImageAEOT[31] applied autoencoders[32–35] for latent embeddings prior to OT from nuclear and chromatin imaging data in breast cancer progression. We believe a similar technique for scRNA-seq data will help us study longitudinal transcriptome dynamics in diseases such as AD. Together, while techniques exist to study spatial and temporal patterns from scRNA-seq data, especially in developmental contexts, the field currently lacks a unified and biologically-interpretable framework to model spatio-temporal dynamics in disease.

Here, to address many of the limitations in existing methods, we introduce sc4D, a biologically-interpretable spatio-temporal (4D) analysis framework for single-cell transcriptomics in disease studies. sc4D applies spatial local-neighborhood analysis of three-dimensional (3D) tissue, autoencoder-based embedded OT for trajectory inference, and *in silico* transcriptome perturbations. Our sc4D framework enables interpolation of intermediate disease states, nomination of key gene markers and reactive cell subtypes, and interpretable cellular perturbations in transcriptome space through latent decoding. We hypothesized that our integrative spatio-temporal approach will reveal critical cell states and nonlinear transcriptome dynamics during disease progression that are not detectable by spatial- or longitudinal-only analyses. We apply sc4D to study the localized cellular response to Aβ plaque and PHF+ tau tangles and longitudinal cellular lineages in AD. Reanalyzing four time-labeled single-cell spatial transcriptomics 3D coronal sections from TauPS2APP mouse models and non-transgenic controls[36], we report critical glial and transcriptome trajectories in AD associated with both the spatial organization of pathological aggregates and disease progression. Notably, we identify activated microglia and astrocyte states that form glial syncytia in response to Aβ plaque. We further nominate transcription factors and spatially enriched gene sets that are predicted to toggle glial activation and regulate inflammatory response to AD. Overall, sc4D readily nominates complex, nonlinear cellular and transcriptome responses to pathologies, guiding therapeutic strategies targeting glia and other cell populations.

## Results

### Study overview

We developed sc4D, a biologically-interpretable spatio-temporal (4D) analysis framework, to investigate the spatio-temporal dynamics of cellular response to pathology during disease progression at single-cell resolution (**Fig 1, Methods**). We analyze longitudinal spatial transcriptomics and pathological imaging datasets that capture spatio-temporal dynamics at distinct time points (**Fig 1a**). First, through local-neighborhood analysis of cells surrounding pathological features from 3D tissue cross-sections, we reveal spatially enriched cell and gene markers (**Fig 1b**). Then, through embedded optimal transport (**Methods**), we reveal shifts in cell state and transcriptome dynamics during disease progression (**Fig 1c**). Finally, by perturbing spatial gene marker expression, we examine the sensitivity of cell state identity to spatial organization near pathology (**Fig 1d**).

**Fig 1.**
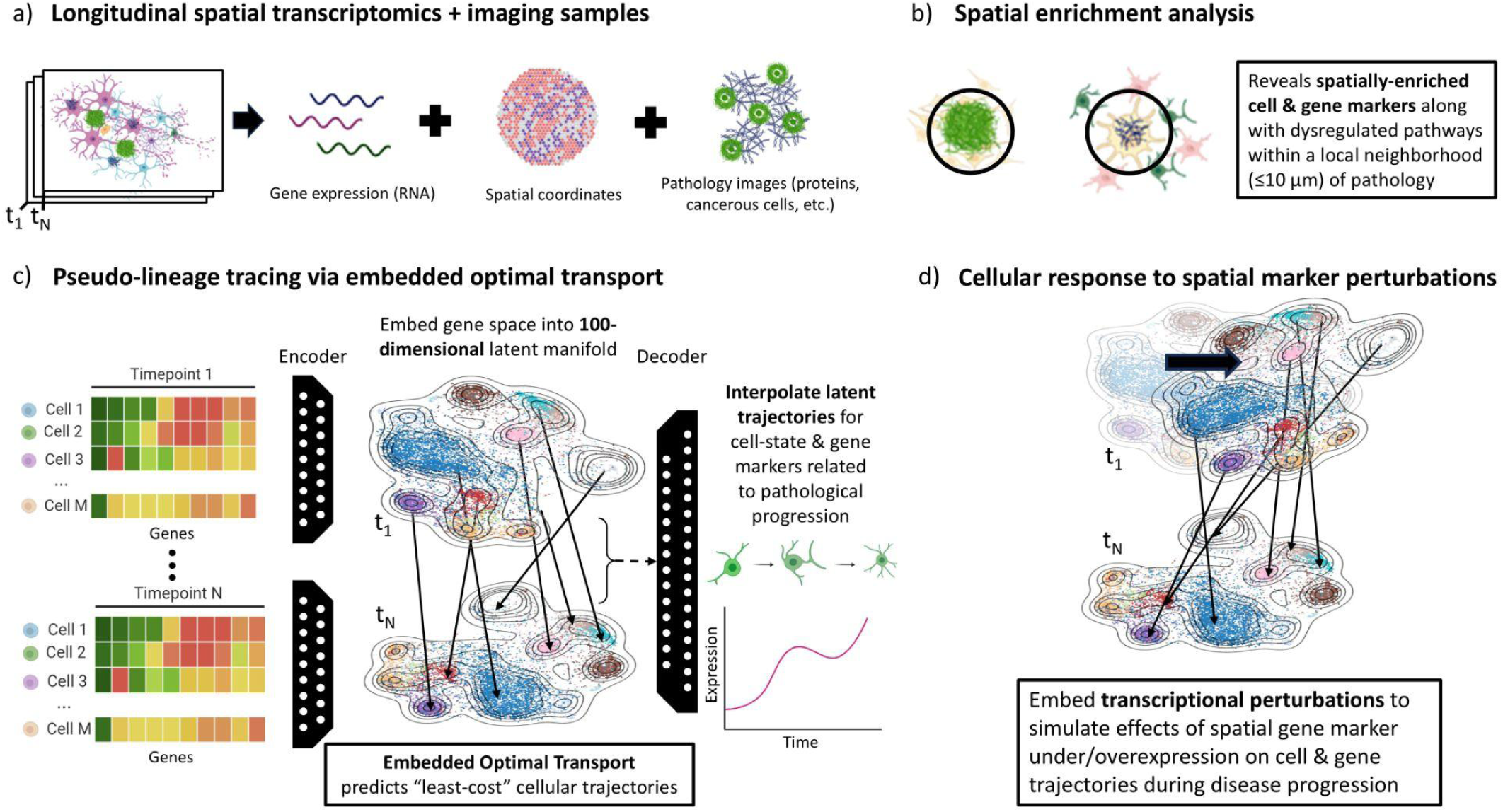
Schematic of our single-cell spatio-temporal analysis (sc4D) framework. We analyze (**a**) longitudinal spatial transcriptomics samples and pathological imaging from different stages of disease progression (e.g., moderate to late stage dementia). (**b**) We identify spatially-enriched cell subtypes, gene markers, and biological processes within a local neighborhood (10-micron radius) of pathological aggregates. (**c**) Using an autoencoder and optimal transport architecture, we embed our cells into a low-dimensional latent manifold to predict cellular differentiation trajectories. We interpolate these trajectories and decode them to gene space to identify key temporal markers of AD progression. Finally, (**d**) we demonstrate the potential of sc4D to enable *in-silico* predictions of cellular and gene expression response to transcriptomic perturbations. Specifically, we perturb spatially-enriched genes prior to embedding to predict their influence on cell and transcriptome trajectories.

In our initial application of sc4D, we analyzed spatial single-cell transcriptomics and imaging-based protein detection data from AD+ and control coronal mouse brain datasets at 8 and 13 months of age with a total of 72,165 cells and 2766 genes across four samples (**Fig S1**)[36]. Spatial mapping (**Fig S1a**) of flattened 3D brain tissue cross-sections revealed the localization of neural and glial cells in heterogeneous brain regions (**Fig S1b**). In addition, we observe a 47% decrease in excitatory neurons and an 86% increase in microglia in AD samples, highlighting neurodegeneration and immune response during disease progression (**Fig S1c, Table S1**). Protein detection data identified Aβ plaque concentrated in the cortical region, spreading into the hippocampus at later stages. Paired helical filament (PHF+) tau was localized to white matter and similarly spread into the hippocampus (**Fig S1d, Fig S2**). Both pathological aggregates were quantified but remained below detection thresholds in nontransgenic controls. Overall, the AD samples display spatio-temporal patterns in cell types, Aβ plaques, and PHF+ tau, aligning with prior studies[37,38]. This suggests that a comprehensive understanding of AD may require joint spatial and temporal analyses to capture its complex cellular and transcriptomic dynamics, motivating the application of sc4D.

### Spatial enrichment analysis reveals markers surrounding Aβ plaque and PHF+ tau tangles

To validate our sc4D approach against known AD biology, we investigated the local neighborhood of a 10 µm radius surrounding Aβ plaque and PHF+ tau (**Fig 2a, Methods**) and identified distinct spatially enriched cell type composition (**Table S3)**, over- and under-expressed differentially expressed genes (DEGs) (**Table S4)**, and corresponding biological pathways (**Table S5**). First, surrounding plaque, we report enrichment of an immune response by the Micro3 subtype (**Fig 2b**) and overexpression of DEGs (**Fig 2c, Table S4),** including *Ctsl*, *Gpnmb*, and *Grn*, involved in over-inflammation[39,40], increased autophagy[41,42], and protective measures in AD[43]. Plaque DEGs were significantly skewed toward overexpression, though we also observed underexpression of neurodevelopment-related genes such as *Cyfip2[44]*. We further report spatial DEG expression to be highly colocalized to plaque aggregates, and to increase in enrichment surrounding plaque during disease progression (**Fig 2e**). Second, surrounding tau, we report enrichment of CA1 excitatory neurons and the Oligo1 subtype (**Fig 2b**) alongside overexpression of DEGs (**Fig 2d, Table S4),** including *Serpina3n* and *Mal*, involved in myelination[45], tau-seed propagation, and neurotoxicity[46–48]. We report a parallel set of underexpressed DEGs, such as *Bsn*, indicating disrupted cytoskeleton development and neuronal function[49] near tau. Between 8- and 13-month samples, the magnitude of z-fold change in spatial DEG expression around tau was minimal compared to plaque, indicating that cells near tau are less broadly reactive to tauopathy than to plaque (**Fig 2e**). Our sc4D approach elucidates the spatial specificity of cell states and gene markers surrounding Aβ plaque and PHF+ tau. These results align with findings from prior scRNA-seq studies of disease-associated cell subtypes[50–52] and gene markers[39–43,45–48], validating our approach.

**Fig 2.**
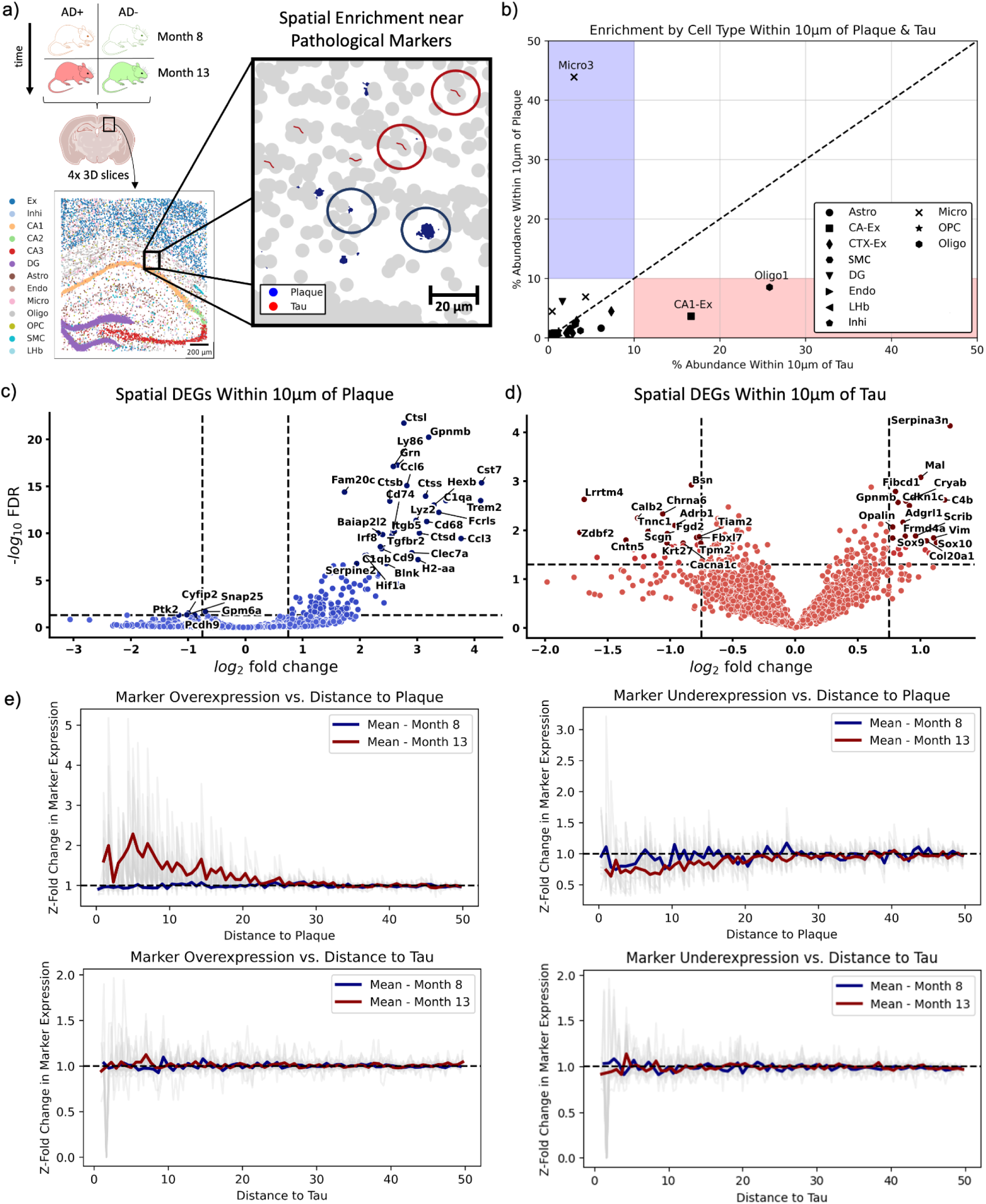
Spatial enrichment analysis within 10 μm of plaque and tau reveals key glial and neuronal subtypes and marker genes. (a) Spatial enrichment analysis of cell types and genes, focusing on the local neighborhood (10-micrometer radius) of Aβ plaque (blue) and tau (red) deposits (**Methods**). Samples were retrieved from coronal slices of AD+ and control mice at 8 and 13 months. Scale bar, 20 μm. (b) Percent abundance of cell subtypes within the local neighborhood of plaque vs. tau. Enriched subtypes display ≥ 10% abundance surrounding plaque or tau. (c-d) Differentially-expressed genes (DEGs) enriched within a 10 micrometer radius of plaque (c) and tau (d) (**Table S4**). We nominate spatially-enriched genes with absolute log2 fold change ≥ 1 and false discovery rate (FDR) ≤ 0.05, with thresholds marked by dashed lines, and label the top 15 over- and underexpressed genes. (e) Over- and underexpression of marker genes relative to distance from plaque and tau pathology (gray), with mean marker expression shown for months 8 and 13.

In addition to confirming established biological processes in AD, sc4D revealed novel spatially enriched biological pathways in the local neighborhood of Aβ plaque and PHF+ tau tangle (**Fig 3, Table S5, Methods**). For example, gene set enrichment analysis (GSEA) surrounding plaque nominated increased macrophage migration (GO:1905517, p=6.70 × 10^−12^), syncytium formation (GO:0000768, p=9.26 × 10^−10^), and antigen processing (GO:0048002, p=3.94 × 10^−9^). GSEA of underexpressed genes surrounding plaque further nominated pathways such as exocytosis of neurotransmitters (GO:0048791, p=2.98 × 10^−3^) and synaptic vesicle recycling (GO:1903423, p=7.70 × 10^−4^), indicating exhaustion of presynaptic machinery near plaque. In the local neighborhood of tau, we nominated increased astrocyte differentiation (GO:0048708, p=3.76 × 10^−7^), glial cell proliferation (GO:0014009, p=1.29 × 10^−3^), and axon ensheathment via oligodendrocytes (GO:0032291, p=2.76 × 10^−3^). GSEA of underexpressed genes surrounding tau further nominated pathways such as regulation of potassium ion transmembrane transport (GO:1901379, p=4.56 × 10^−2^) and protein localization to postsynapse (GO:1904377, p=3.13 × 10^−3^), indicating disruption of ion homeostasis and synaptic organization near tau pathology. Finally, we report a global astrocyte response (GO:0048708) surrounding Aβ plaque (p=1.43 × 10^−2^) and PHF+ tau (p=4.65 × 10^−3^). Together, these spatially enriched terms indicate potential mechanisms of neurodegeneration via loss of synaptic function surrounding tau[12,53] and glial syncytium formation surrounding Aβ plaque. Syncytium formation has been previously characterized between microglia and astrocytes surrounding degenerative neurons[53–55]. Moreover, our results support an enrichment of Aβ plaque formation surrounding tau aggregates through amyloid fibril development, providing spatial specificity to prior studies[56]. Overall, spatially-resolved enrichment analysis via sc4D improves our understanding of AD by nominating attractive hypotheses for disease mechanisms and heterogeneous cellular responses to pathologies (**Discussion**).

**Fig 3.**
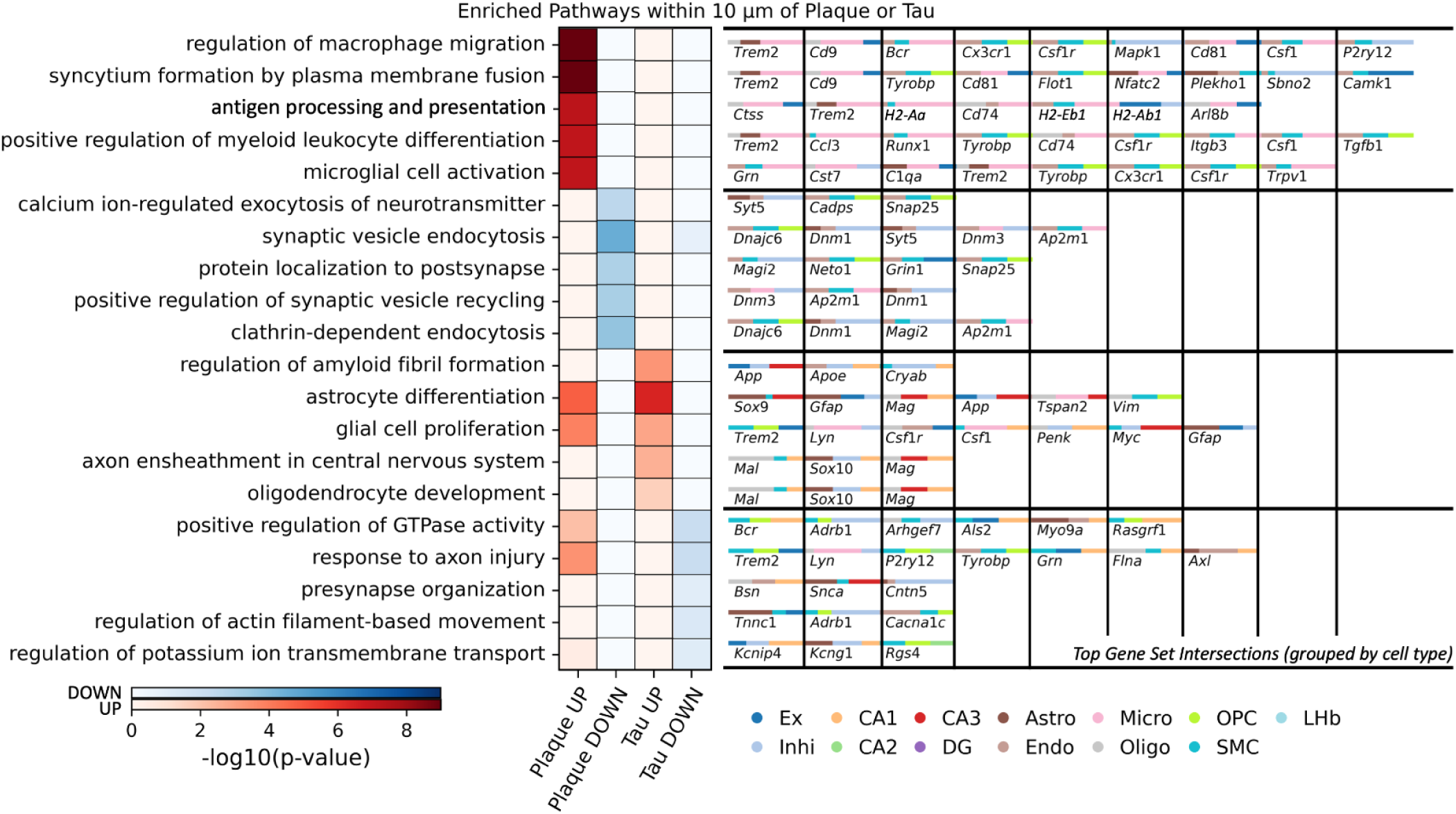
Spatially-enriched pathway analysis reveals syncytium formation in microglia and glial remodeling near tau tangles. Gene set enrichment analysis (GSEA) reveals enriched pathways from Gene Ontology Biological Process surrounding plaque and tau. We report the top five statistically significant pathways for genes overexpressed (UP) and underexpressed (DOWN) surrounding each aggregate as calculated by g:Profiler (**Methods**). We also report the top ten gene set intersections, defined by false discovery rate (FDR)-significant enrichment surrounding plaque and tau, along with the relative enrichment of each cell type in which each intersecting gene is differentially expressed.

### Embedded optimal transport for trajectory inference in Alzheimer’s disease

Alzheimer’s disease typically progresses over seven to ten years following diagnosis[13]. Given this timescale, we hypothesized that, beyond spatial enrichment analysis near plaque and tau, characterizing the progression between middle and late-stage dementia would nominate critical disease markers and mechanisms. To that end, we first compared longitudinal transcriptome samples and identified enriched biological pathways during late-stage dementia across control and disease samples (**Fig 4, Table S6-7, Methods**). In disease samples, pathways such as protein localization to synapse (GO:0035418, p=3.89 × 10^−16^) and postsynaptic membranes (GO:1903539, p=6.16 × 10^−13^) are highly enriched, highlighting alterations in synaptic architecture. Additionally, synaptic vesicle recycling (GO:0036465, p=4.34 × 10^−15^) and endocytosis (GO:0048488, p=1.03 × 10^−12^) are enriched, indicating disruptions in neurotransmitter release. Similarly, we report enrichment of both GABAergic (GO:0051932, p=9.79 × 10^−7^) and glutamatergic transmission (GO:0051966, p=1.52 × 10^−9^) during AD progression, but only enrichment of glutamatergic transmission (GO:0051966, p=2.42 × 10^−4^) in control samples. Lastly, we found dysregulation of dendritic spine development (GO:0060998, p=8.82 × 10^−12^) and morphogenesis (GO:0061001, p=1.33 × 10^−9^). These findings indicate that alterations in synaptic architecture, dendritic spine development, and neurotransmitter release are central to AD progression. In addition, differential enrichment in GABAergic and glutamatergic neurotransmission between disease and control samples indicates dysregulation of inhibitory-excitatory balance in AD. The observed longitudinal dynamics in pathway enrichment motivated us to pursue cell type-resolved modeling of disease progression in AD.

**Fig 4.**
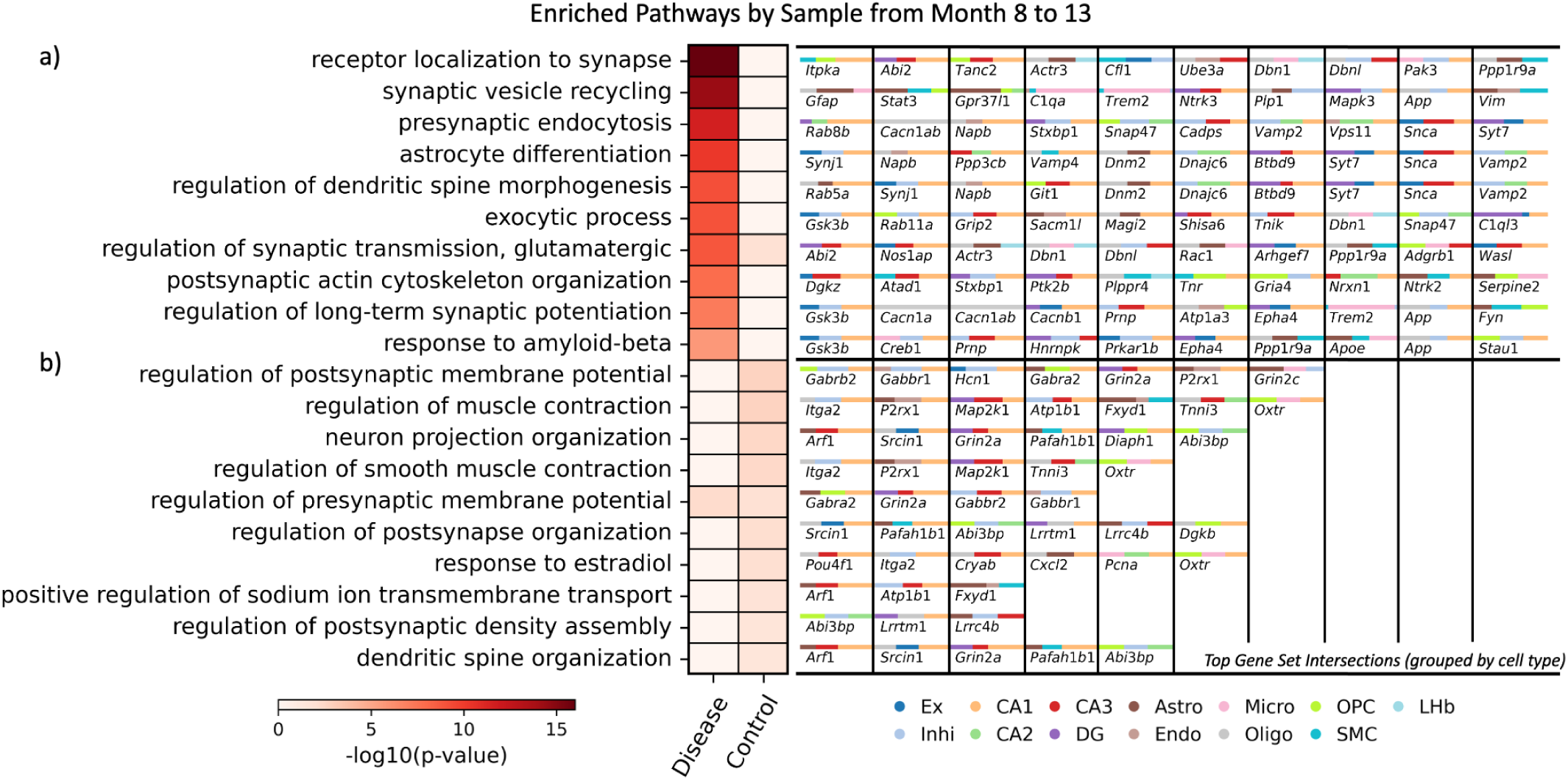
Temporally-enriched pathway analysis between 8- and 13-month samples identifies dysregulated synaptic transmission during AD progression. We report the top 10 pathways from the Gene Ontology Biological Process for control and disease samples as calculated by g:Profiler (**Methods**) and rank statistical significance by log-scaled g:SCS ontology-adjusted p-value. We also report the top ten gene set intersections, defined by false discovery rate (FDR)–significant enrichment between 8- and 13-month samples, along with the relative enrichment of each cell type in which each intersecting gene is differentially expressed.

To gain a high temporal-resolution understanding of cellular states and transcriptome dynamics during AD progression, we investigated longitudinal dynamics of cellular trajectories via sc4D by applying optimal transport (OT) to lower-dimensional embeddings (**Methods**). Specifically, we constructed a 100-dimensional latent embedding of the time-labeled single-cell transcriptomes through a trained autoencoder model (**Fig 5a, Fig S3-S5**, **Methods**). As a baseline, we also randomly sampled from a fitted zero-inflated Poisson distribution, and validated that our autoencoder model converged on substantially smaller reconstruction loss (**Fig S4, Methods**).

**Fig 5.**
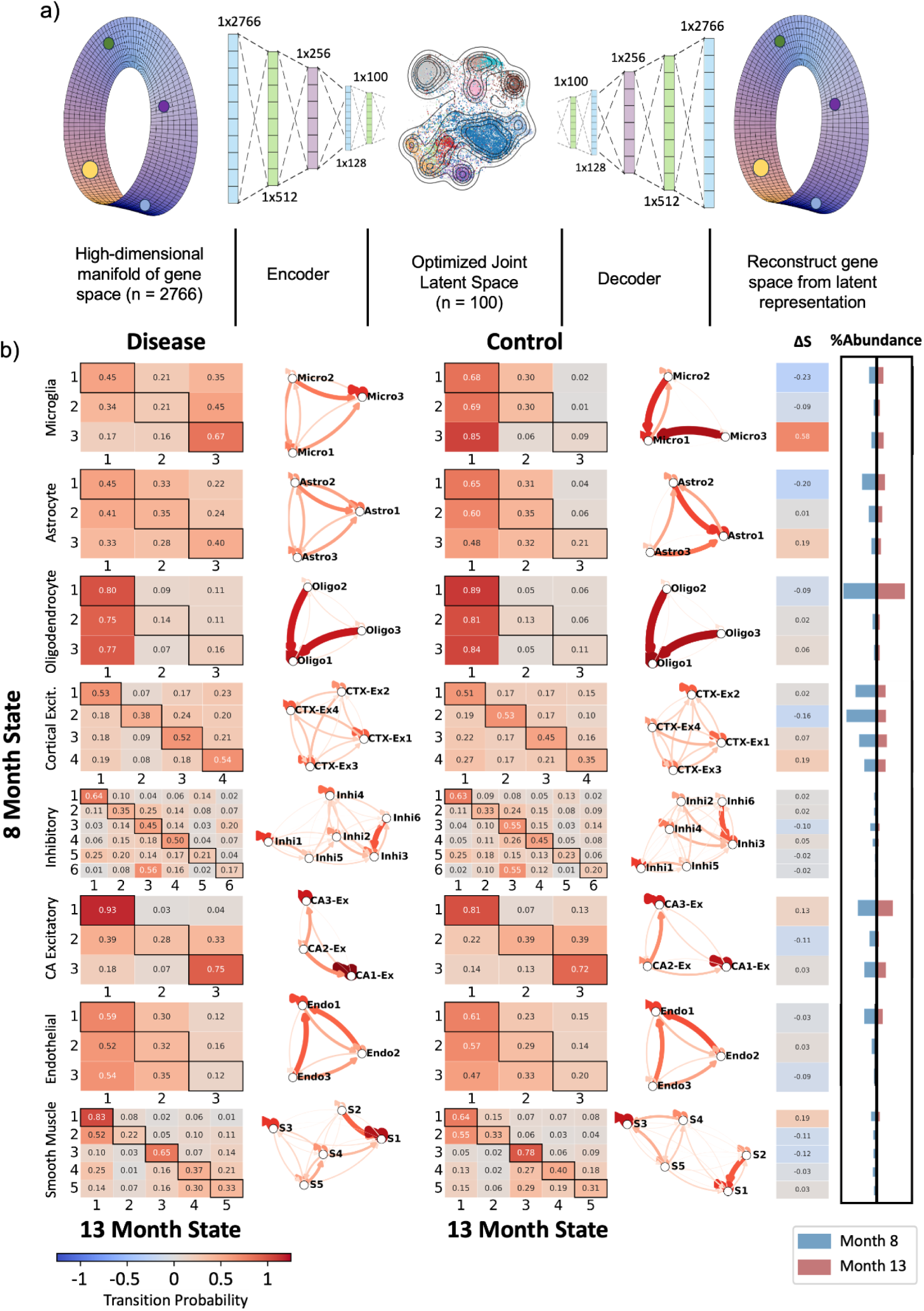
Single-cell autoencoder enables embeddings to predict cell state transitions and spatial marker expression over time through optimal transport. **(a)** Schematic of applying an autoencoder for scRNA-seq dimensionality reduction and reconstruction. **(b)** The transition matrices display the transition probability between cell states from months 8 to 13 in disease and control samples (**Fig S4**), as predicted by optimal transport (OT) (**Methods**). For intuition, we also visualize this transition matrix as a directed graph for each cell type. For each cell state, we report the stability metric (**Methods**) and the differential stability (ΔS), defined as the difference in stability between disease and control samples. We also plot the percent composition of each cell subtype at 8 and 13 months in our AD+ samples.

Given distinct longitudinal shifts in functional pathways, we applied trajectory inference in sc4D via unbalanced OT to our time-labeled latent embeddings to investigate whether these shifts mapped to key cell state transitions (**Methods**). Unbalanced OT efficiently redistributes cells between our 8- and 13-month samples (**Fig S5**) by allowing cells in the smaller set to couple with multiple cells in the larger set, facilitating cellular proliferation and reconstructing trajectories despite heterogeneous samples. In order to evaluate how cell states respond to disease stressors, we defined stability from the transition probability matrices as the self-transition probability, nominating cell states which conserve their identity during disease progression as more stable (**Methods**). We observed a gain in stability across several glial states between disease and control samples (**Fig 5b**), which we interpret as shifts toward stress-responsive states and immune-glial reactivity in the vicinity of plaque and tau[57]. Based on the transition probability between states for each cell type from our OT mappings (**Fig 5b**), we primarily find transitions along the main diagonal, indicating most cell populations maintain their identity during AD progression. Exceptions to this pattern in conjunction with gain of stability indicated shifts toward states likely correlated with disease and stress response[58]. We report transitions toward Micro3 and Astro3 subtypes, both of which show gain of stability. Importantly, stability changes are not always mirrored by changes in cell abundance: glial states such as Micro3 and Astro3 exhibit gain of stability even without proportional changes in abundance, indicating that OT captures reprogramming of cellular states beyond population size dynamics. Through optimal transport-based trajectory inference, we find shifts toward key astrocyte and microglia states during AD progression, with cell stability linking stress-induced glial reactivity to the glial syncytium hypothesis from our spatial analysis (**Discussion**).

To assess the ability of embedded OT to recover transcriptomic shifts correlated with disease response, we interpolated trajectories and tracked spatial DEG expression during AD progression per cell state, specifically focusing on trends in neuronal cell types (**Methods, Fig 6**). Neuronal expression near amyloid plaques shows a non-directional reduction of both Plaque (UP) and Plaque (DOWN) gene sets, whereas neurons surrounding tau follow the expected directional trends, i.e. increased expression of Tau (UP) genes, decreased expression of Tau (DOWN) genes (**Fig 6a**). We interpret this difference as reflecting the distinct mechanisms of pathology, in which plaques create a hostile microenvironment leading to neuronal loss (**Fig 3**), producing net reductions in expression across gene sets. In contrast, tau pathology is more neuron-intrinsic and elicits directed transcriptional programs, e.g., induction of stress/repair genes and repression of synaptic genes in the affected neurons[59]. Together, these results indicate that plaque primarily drives neuronal loss, while tau drives neuron-intrinsic transcriptional remodeling during disease progression.

**Fig 6.**
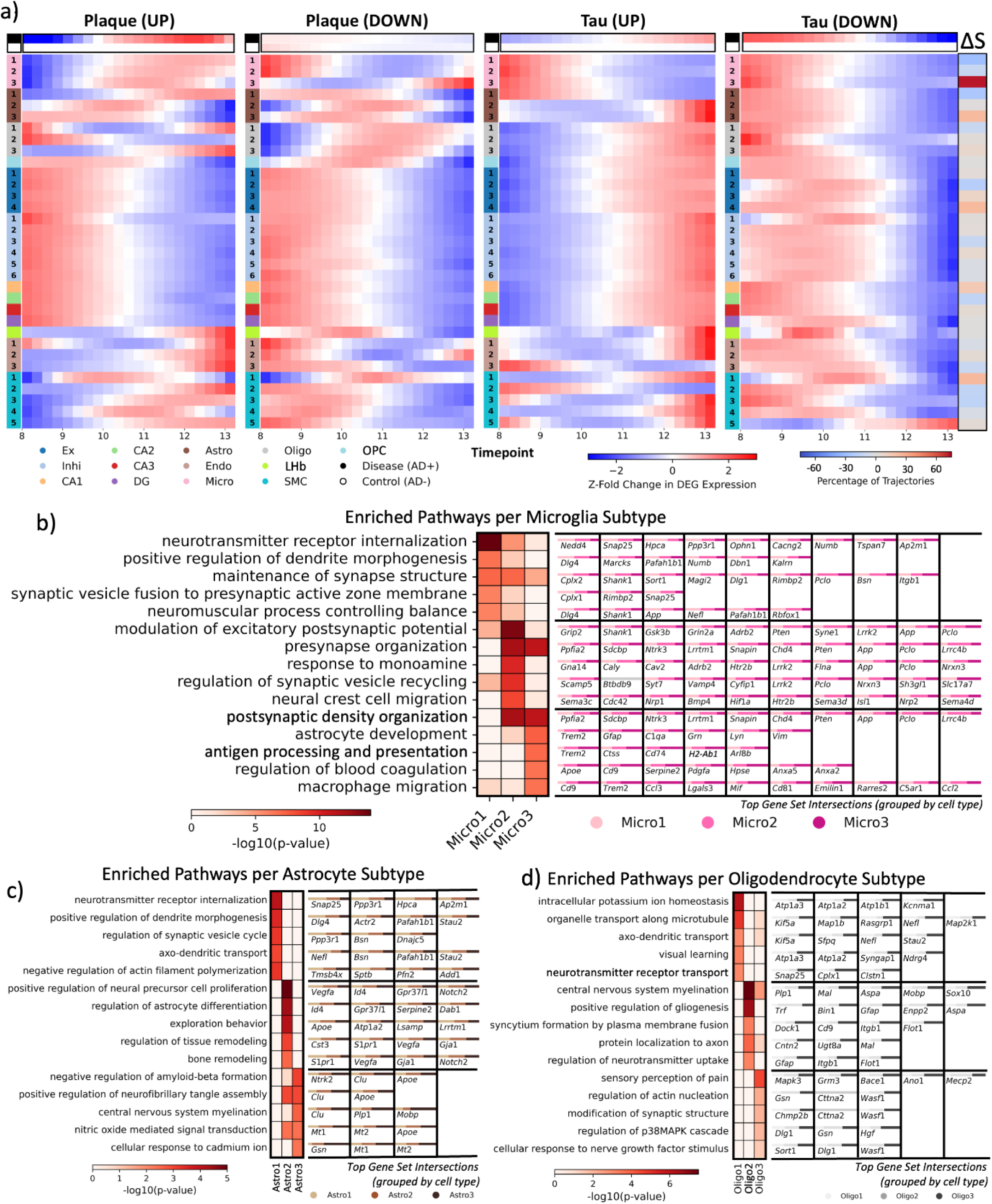
Longitudinal patterns of spatially-enriched gene sets and subtype-resolved pathway analysis reveal reactive microglia and astrocyte states involved in amyloid-tau pathology. **(a)** We report the z-fold change in mean expression of the four gene sets defined in Fig 2e over interpolated OT paths from 8 to 13 months. We compare the mean expression of the genes across disease vs. control samples and by cell state. The top five enriched pathways as calculated by g:Profiler and rank statistical significance by log-scaled g:SCS ontology-adjusted p-value for each (**b**) microglia, (**c**) astrocyte, and (**d**) oligodendrocyte subtype (**Methods**). We also report the top gene set intersections, defined by false discovery rate (FDR)–significant enrichment of each state relative to all other cell states, along with the relative enrichment of each cell state in which intersecting genes are differentially expressed.

In glial populations, spatial DEGs associated with plaques, Plaque (UP) and Plaque (DOWN), are expressed earliest in the most reactive cell states and progressively later in intermediate and homeostatic states (**Fig 6a**). Microglia, for example, show strongest early expression and highest stability in Micro3, which is characterized by dysregulated blood coagulation (GO:0030193, p=2.51×10^−7^) downstream of neuroinflammation[60,61], microglial migration (GO:1905517, p=2.73×10^−7^), and *Apoe/Trem2*-driven programs (**Fig 6b, Table S9-10**), aligning with the disease-associated microglia phenotype[60]. Micro2 and Micro1 show successively delayed expression, consistent with their more homeostatic phenotypes maintaining synaptic structure (GO:0099558, p=8.76×10^−8^) and vesicle recycling (GO:1903421, p=4.71×10^−10^). Astrocytes (**Fig 6c**) and oligodendrocytes (**Fig 6d**) follow the same pattern: Astro3 and Oligo2, enriched for amyloid (GO:1902430, p=8.60×10^−4^) and tau (GO:1902998, p=1.09×10^−3^) stress responses, gliogenesis (GO:0014015, p=2.83×10^−7^), increased myelination (GO:0022010, p=3.97×10^−8^), and syncytium formation (GO:0000768, p=5.59×10^−4^), display the earliest plaque-associated transcriptional induction, whereas their less reactive counterparts (Astro2/1, Oligo3/1) respond later (**Table S9-10**). Across glial lineages, the timing of spatial DEG induction mirrors both the functional phenotype and stability of each cell state, highlighting reactive glial states with the highest stability (Micro3, Astro3, and Oligo2) as the earliest and most plastic responders to plaque-associated stressors, with spatially-enriched transcriptional programs progressively propagating into less activated populations.

Overall, we report longitudinal patterns in transcriptome profiles and spatial DEGs, with joint neuronal expression of spatial DEGs, and a staggered glial response to Aβ plaque. As such, these spatial DEGs likely drive disease-associated transcriptional remodeling in neurons and glial activation, serving as key mediators of the cellular response to plaques.

### Modeling *in-silico* drug perturbations with embedded OT

To study the relationship between longitudinal cell lineages and spatial DEG expression, we modeled four *in silico* perturbations to latent embeddings of transcriptomes with sc4D (**Fig S6**). Specifically, we assess how plaque or tau influences cell identity changes along the disease trajectory by emulating localized pathology removal through over- or under-expression of spatially enriched genes surrounding plaque and tau regions (**Methods, Fig. 7a–b, Table S11**). Perturbed transition matrices across all gene sets and supporting independent experimental findings (**Fig 7c-d**) reported microglial shifts away from activated Micro3 toward homeostatic Micro1/2, which simultaneously lost stability, indicating that Micro3 activation is tightly linked to the presence of plaques and tau. Astrocytes showed a similar pattern, with transitions from reactive Astro3 toward homeostatic Astro1 via Plaque (DOWN) perturbation. In contrast, oligodendrocytes shifted toward the activated Oligo2 state, particularly through the Oligo3 to Oligo2 pathway via Tau (DOWN) perturbation. This counterintuitive response may reflect a compensatory mechanism: homeostatic Oligo1 are already located near tau (**Fig 2b**), and removal of localized tau pathology signals may relieve inhibitory cues, enabling cells to activate neuro-glial signaling and myelination programs associated with Oligo2 (**Fig 6d**). Out of the four spatial DEG perturbations, Plaque (DOWN) perturbation, which is most closely linked to CAMTA1 activation, is predicted to most broadly induce homeostatic states (namely, Micro1, Astro1, and Oligo1). From this result, we nominate Plaque (DOWN) genes (e.g., *Gpm6a, Cyfip2, Snap25, Ptk2*) as likely glial activators in AD which display both spatial and longitudinal enrichment. Overall, perturbation modeling demonstrates that both pathology and therapeutic intervention can reshape glial cell states, highlighting the central role of spatially enriched DEGs in driving glial identity, activation, and plasticity during disease progression (**Discussion**).

**Fig 7.**
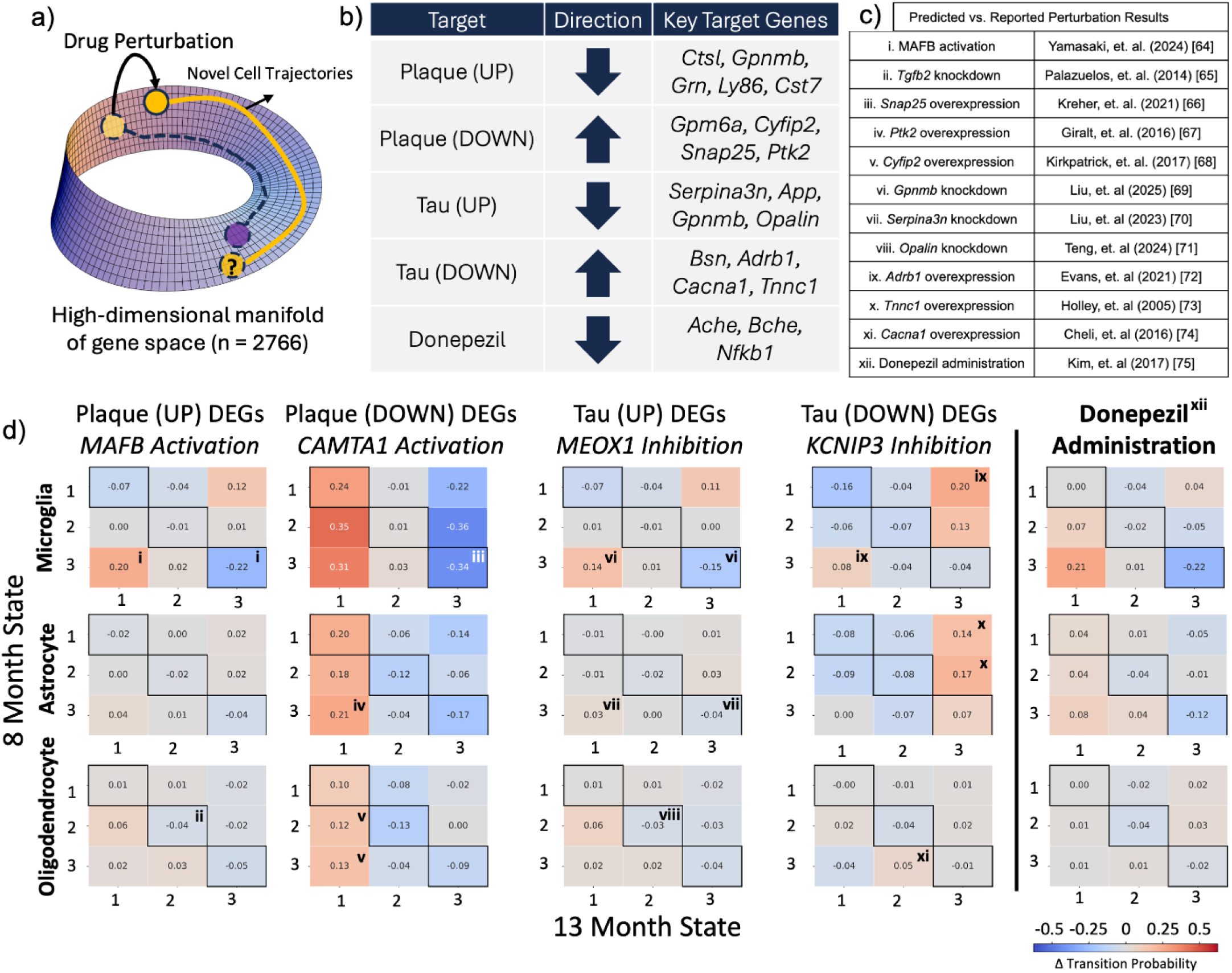
Modeling unseen perturbations on single-cell samples based on DEGs and known pharmaceutical profiles reverses glial activation. (**a**) Schematic demonstrating the effect of unseen drug-induced perturbations in the high-dimensional gene space (n_genes_ = 2766) of a cell and the expected changes to optimal transport-predicted cell state trajectories. (**b**) Five *in-silico* gene perturbations were individually applied to our AD+ single-cell samples before trajectory inference based on DEGs and known pharmacogenetic profiles (**Methods**). (**c**) Independent experimental findings supporting OT-predicted perturbation results. (**d**) Net transition matrices and differential stability of each cell state are shown across all perturbations, along with cross-references to independent experimental findings supporting perturbation results (**Table 1**). Enriched transcription factors regulating each gene set are also noted (**Methods**).

Beyond modeling the effects of spatially enriched markers, we explored cell-type-resolved transcriptomic changes resulting from pharmaceutical interventions. Specifically, we modeled the impact of Donepezil administration, an acetylcholinesterase (ACHE) inhibitor and approved AD therapy[62]. Perturbation analysis showed a shift of microglia and astrocytes toward homeostatic states. This transition supports neuronal function by reducing inflammation and strengthening neurotransmission (**Fig 6d**), consistent with the clinical benefits of Donepezil in improving cognitive performance across mild, moderate, and severe dementia[63]. These results illustrate how the sc4D framework can capture transcriptomic and pharmacogenetic perturbations to resolve cell state trajectories, providing a path to design candidate interventions that bias glial and neuronal identity toward phenotypes most protective against disease progression.

## Discussion

Here, we introduce sc4D, a biologically-interpretable spatio-temporal (4D) analysis framework for single-cell transcriptomics in disease studies that integrates spatial transcriptomics with trajectory inference and *in silico* perturbations through embedded optimal transport (OT). As a proof of concept, we present sc4D’s application to a longitudinal spatial single-cell transcriptomics dataset of Alzheimer’s disease (AD) to capture complex cellular and transcriptome dynamics in disease response from 3D tissue cross-sections. Our framework identifies a joint and complex glial response of microglia, astrocytes, and oligodendrocytes to plaque and tau during disease progression. sc4D offers attractive hypotheses through a more comprehensive view of dynamic cellular responses in disease that traditional spatial- or temporal-only analysis pipelines cannot detect.

Computationally, we demonstrate several advantages of sc4D’s embedded OT technique for trajectory inference from scRNA-seq samples over existing approaches[76,77]. First, it enables modeling of cellular differentiation over chronic disease timescales (e.g., months to years) by integrating time-series data into inferred trajectories[31], addressing limitations of existing methods effective only over hours or days[76,77], such as in developmental contexts. Second, it interpolates intermediate transcriptomes from time-labeled scRNA-seq data, potentially reducing the need for high-resolution time-series datasets. Further studies are necessary to validate whether this approach, with temporal data sufficiently representative of potential cell states, can reveal intricate cellular and transcriptomic dynamics from sparse longitudinal data. Third, it employs autoencoder embeddings, which are known to outperform traditional dimensionality reduction methods like t-SNE in improving interpretability[78]. Further, autoencoder embeddings capture nonlinear dynamics more accurately than techniques like PCA[79], which make unrealistic assumptions that linear distances in transcriptome space are representative of cell similarity. As a result, application of autoencoders as in sc4D enables more representative trajectory inference through embeddings that reflect complex biological patterns. Fourth, it models downstream effects of transcriptome perturbations, improving biological intepretability through the autoencoder model by identifying key drivers of cellular differentiation and uncovering potential therapeutic targets to influence cell lineages in disease contexts. This could enable therapeutic control over processes such as immune response, allowing precise modulation of how cells react to disease—for example, reducing neuroinflammation while still enabling effective clearance of pathology. Overall, sc4D captures nonlinear cellular and transcriptomic dynamics in disease contexts while elucidating the effects of transcriptome perturbations on cell lineages. In combination with existing spatial transcriptomics analysis at each time point, our results underscore the benefit of jointly considering spatial organization and longitudinal progression in characterizing cellular response to localized diseases.

Biologically, sc4D revealed spatio-temporal enrichment of oligodendrocyte and activated microglial subpopulations, syncytium formation, astrocyte activation, synaptic dysfunction, and increased myelination in proximity to Aβ plaque and PHF+ tau tangles. Longitudinal analysis reveals similar differentiation toward activated microglial and astrocyte populations, along with localized increases in spatially enriched marker expression in glial populations during AD progression. Together, these results provide spatial specificity to the key role of activated microglia[60] that react in proximity to Aβ plaque, increased myelination by oligodendrocytes surrounding tau-positive neurons, and global astrocyte activation as key components of cellular response to AD. Furthermore, we hypothesize that microglial and astrocyte lineages toward activated subtypes, along with syncytium formation surrounding plaque, provide evidence of glial syncytia that envelop and compress Aβ plaque aggregates[80], supported by recent studies that identified these syncytia to support synaptic health and long-term potentiation[53,54]. These syncytia may serve to isolate the toxicity of Aβ plaque and support neuronal function in the presence of plaque. Finally, perturbations to spatially-enriched marker expression via sc4D are predicted to limit microglial and astrocyte activation, indicating a correlation between proximity to Aβ plaque or PHF+ tau and activated glial identity.

From this initial study of sc4D for spatio-temporal single-cell analysis in disease contexts, we recognize five future directions to extend upon our technique. First, we applied trajectory inference between only two time points. Analyzing higher resolution temporal data would result in more accurate and likely more complex lineages during disease progression. Second, we may incorporate multi-omics data into our autoencoder embeddings[81], such as integrating both chromatin imaging and scRNA-seq data, to reveal intricate mechanisms of cellular response. We can also incorporate spatial coordinates to inform our OT trajectories via Fused Gromov-Wasserstein OT[82], allowing us to jointly study how spatial organization and transcriptome profiles impact cell lineages. Third, while we apply perturbations based on predicted pharmacogenetic profiles, future studies may utilize cell type-specific data[83] to reveal key perturbagens and predict downstream therapeutic effects on cell lineages. Fourth, while this application of sc4D focused on Alzheimer’s disease, it can also be used to study cellular and transcriptome responses to other localized diseases, such as spinal cord injuries[17] and in tumor microenvironments[84]. Finally, future studies can implement sc4D to model cellular dynamics from early to late-stage disease from human datasets, for example, enhancing disease forecasting and early interventions.

Overall, our results demonstrates the importance of a unified single-cell framework that integrates spatial organization and longitudinal dynamics to offer a holistic view of disease response.

## Methods

### Spatio-temporal single-cell and protein marker expression dataset for AD

To investigate spatio-temporal molecular changes in AD progression, we reanalyzed spatially resolved single-cell RNA sequencing and imaging of two AD protein markers, Aβ plaque and PHF+ tau, from 3D coronal mouse brain sections[36]. We profiled coronal cross-sections with a voxel size of 95 × 95 × 350 nm from two AD-transgenic (AD+) and two control non-transgenic mice at 8 and 13 months. We used the TauPS2APP model, a well-established triple-transgenic AD model[14] that expresses the P301L mutant of human tau and PS2N141I and APPswe (PS2APPhomo; P301Lhemi). This model develops moderate dementia at 8 months and late-stage dementia at 13 months in cortical and hippocampal regions. Using STARmap PLUS[36], we simultaneously retrieved gene expression values from 72,165 cells across 2,766 genes, spatial coordinates, and locations of Aβ plaque and PHF+ tau aggregates from intact 3D coronal tissue sections. We filtered, normalized, and scaled all expression data as described in[36]. For cell annotation, we applied 33 cell types and sub-cluster labels from the dataset and listed the cell counts per sample (**Table S1**). We confirmed the specificity of these labels using key marker genes identified with the ‘rank_genes_groups’ method in Scanpy[85] (**Table S2**).

### Spatially-resolved visualization of cells and AD protein markers

We visualized the spatial organization of cells with cell type or region annotations through the ‘spatial_scatter’ method implemented in the Squidpy Python package[86]. For each sample, we counted the number of cells and reported the composition of cell populations. We overlaid Aβ plaque and PHF+ tau sites from binarized images through the ‘imshow’ method in Matplotlib[87].

### Aβ plaque and PHF+ tau-associated spatio-temporal enrichment analysis

We conducted spatio-temporal enrichment analysis of cell subtypes and gene expression in AD based on the spatial organization of Aβ plaque and PHF+ tau, along with spatial scRNA-seq profiles from moderate and late-stage AD mice datasets (**Fig S1**).

We nominated spatially-enriched subtypes by evaluating the percent abundance of each within the local neighborhood (10 micrometers) of Aβ plaque[88] or PHF+ tau in our two AD+ samples (**Fig S2**). We nominated cell subtypes that exhibited ≥ 10% abundance within the local neighborhood of protein markers as plaque- or tau-enriched.

To identify spatially differentially-expressed genes (DEGs) associated with plaque and tau, we compared the expression profiles of cells within the local neighborhood of plaque or tau to all other cells. We calculated each gene’s log-fold change, dispersion, variance, and p-value via the Wald test in PyDESeq2 [89]. We defined DEGs as those with a false discovery rate (FDR) ≤ 5% and an absolute log-fold change > 1, which we visualize as a volcano plot. We define four gene sets, Plaque (UP), Plaque (DOWN), Tau (UP), and Tau (DOWN), based on whether DEGs were over- or underexpressed surrounding plaque and tau. We visualize spatial enrichment by the z-fold change in marker expression for each gene set surrounding plaque and tau from 0-50 μm radius in our 8 and 13-month samples. We then identified differentially regulated biological pathways surrounding each protein aggregate via gene set enrichment analysis (GSEA) of our plaque- and tau-associated spatial DEGs. Specifically, we conducted an over-representation analysis with the Gene Ontology Biological Processes set[90,91] via the g:GOSt functional profiling method in g:Profiler[92] with g:SCS ontology-corrected p-value. We filtered for specific functional pathways by limiting the term size to a maximum of 100 genes and setting the background to all genes measured in our dataset. We visualized the top five selected enriched pathways for each of the over- and underexpressed plaque- and tau gene sets, ranked by -log_10_(corrected p-value), as a heatmap through Seaborn[93]. To determine cell type specificity within each functional pathway, we also display the top ten gene set intersections, defined by false discovery rate (FDR)-significant enrichment surrounding plaque and tau, together with the relative significance of each cell type based on the distribution of absolute log-fold changes for the intersecting genes.

To investigate AD-specific changes during the progression from moderate to late-stage dementia, we nominated temporally enriched DEGs between stages across control samples, AD+ samples, and all samples through the Wilcoxon rank-sum test in Scanpy’s ‘rank_genes_groups’ method. Following previously defined parameters, we performed GSEA via g:Profiler on our temporally enriched DEGs. We visualized the top ten biological pathways, ranked by -log_10_(corrected p-value), for control samples and AD+ samples as a heatmap, along with the top ten gene set intersections as described previously.

### Non-linear single-cell expression embeddings through autoencoders

To prepare for model training, we combined normalized scRNA-seq count data across all four samples. To maintain a larger validation and test dataset for more robust results, we implemented a 60-20-20 split between training, validation, and held-out test sets across all cells.

To reduce the dimensionality of scRNA-seq data, we applied an autoencoder model for 100-dimensional single-cell embedding through TensorFlow’s sequential API[94]. We designed an autoencoder with an input of 2766 genes, featuring four dense layers using the Rectified Linear Unit (ReLU) activation function and a 100-dimensional bottleneck layer (**Fig S3**). We employed the Adam optimizer[95] with a learning rate initialized at 1e-3 and a mean squared error (MSE) loss function, and trained the model for 500 epochs with a batch size of 256. The close agreement between validation and training losses indicates that the model generalizes well to unseen data (**Fig S4**).

As a quality control for autoencoder training, we compared the autoencoder’s training and validation loss to the MSE loss from 500 expression matrices (one per training batch) randomly sampled from a zero-inflated Poisson model fitted to our normalized scRNA-seq count data through StatsModels Formula[96]. We calculated the average MSE loss between random samples generated from a zero-inflated Poisson model and the held-out test data, providing a baseline for comparison with the autoencoder’s performance.

To reduce the dimensionality of our single-cell samples and visualize the data, we first projected the single-cell expression data to a 100-dimensional latent space using the trained encoder. A linear shift in the latent space corresponds to a complex nonlinear shift in the original high-dimensional gene expression vector.

To visualize and quantify the shift in the distribution of cell populations in latent space between 8 and 13 months, we applied a kernel density estimation (KDE) topographical overlay and applied the Kolmogorov-Smirnov (KS) test. We first apply a KDE overlay with six contour levels via Seaborn[93] onto a two-dimensional Uniform Manifold Approximation and Projection (UMAP)[97] of our single-cell samples, with Euclidean distance and 5 nearest neighbors to preserve local resolution. Second, we visualized the count distribution of each cell type in a latent dimension (UMAP2), which is smoothed by a Gaussian filter (*σ* = 5) in SciPy[98]. We plotted these smoothed distributions on the y-axis of each UMAP (**Fig S5**). Third, we applied the two-sample Kolmogorov-Smirnov (KS) test via SciPy to quantify shifts between our 8 and 13-month distributions for the control and AD+ samples at the cell-type level across each of our 100 latent dimensions. For each top-level cell type distribution across all 100 dimensions, we reported the most significant KS statistic ranked by Bonferroni-corrected p-value (α = 4.5×10^−5^) (**Table S8**).

### Embedded unbalanced optimal transport for lineage tracing

To determine single-cell trajectories and capture nonlinear relationships in the data, we applied unbalanced optimal transport (OT) on our latent embeddings implemented in the Python Optimal Transport (POT)[99] package.

Specifically, we solved for the unbalanced OT plan, π, between the source (8 months) and the target (13 months) latent embeddings by the entropy-regularized and Kullback-Leibler divergence-parameterized unbalanced OT formulation (**Equation 1**)[31]:

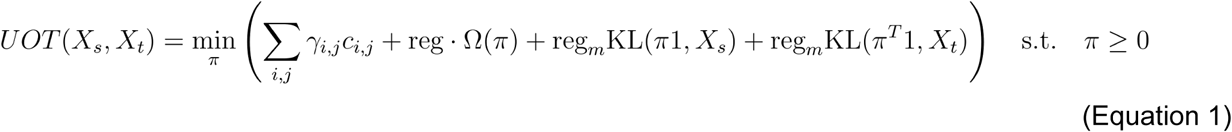

where the first term represents the product of the transport flow between timepoints *γ_i,j_* and cost function *c_i_*_,*j*_, the second term represents entropic regularization, Ω(*π*), and the last two terms represent the marginal constraints via Kullback-Leibler (KL) divergence. For the cost matrix, *c*, we used a normalized Euclidean distance between the source and the target latent embeddings, labeled *X*_s_ and *X*_t_ respectively (**Equation 2**):

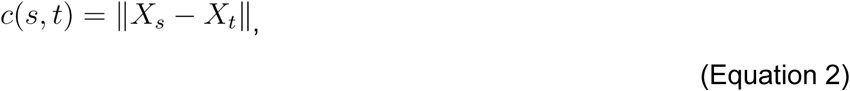

, which we obtained by POT’s ‘dist’ method and subsequently normalized by the maximum distance value so that the Euclidean distance is scaled within the [0, 1] interval. The entropic regularization term encourages a smooth and strictly convex transport plan between the unbalanced distributions, allowing for a distinct mapping between cells and efficient approximation with the Sinkhorn-Knopp algorithm[100]. Lastly, by incorporating the KL divergence parameters, we penalized violations of the marginal constraints and adjusted the regularization parameter *reg_m_* based on the differences between our source and target distributions.

In our application, we used the Sinkhorn-Knopp algorithm implemented in POT’s ‘sinkhorn_unbalanced’ method to approximate our transport plan with *reg* set to 0.005 and *reg_m_* set to 0.05. Low entropic regularization prioritizes minimizing transportation costs over enforcing strict sparsity constraints on the transport plan and allowing multiple predicted lineages per cell. Low KL regularization increases penalties for marginal constraint violations, reducing sensitivity to mass shifts between the source and target distribution.

In order to accurately model mature cell type dynamics in disease contexts, where cellular transdifferentiation is expected to be rare[101], we constrained the unbalanced OT coupling so that mass could only flow between cells assigned to the same annotated cell type and between cell states (e.g., Micro1 to Micro2)[102]. We implemented this by masking the cost matrix by applying explicit zero priors on cellular trajectories between cell types and renormalizing within-type marginals. These constraints preserved biologically plausible within-type maturation and prevented artifactual cross-type mappings while still allowing unbalanced mass transfer and entropic smoothing within states (*reg* = 0.005, *reg_m_* =0.05) as previously described. By constraining transport in this way, we traded sensitivity to rare true transdifferentiation events for greater specificity of within-type lineage inference.

To model cellular proliferation, where target cells at 13 months have one progenitor, we selected lineages from the transport map by selecting the lowest transport cost for each target cell. As a result, source cells at 8 months may have more than one target cell.

### Trajectory inference for cellular and transcriptome change

To visualize cell state dynamics in AD as predicted by OT, we calculated a transition matrix to determine the percentage probability for each cell state to transition between states from 8 to 13 months. We normalized each row of the transition matrix to a sum of 100% probability and generated separate matrices for diseased and control samples. We also visualized the transition matrix as a directed, fully-connected graph, where nodes represent cell states and directed edges are colored by the transition probability, including loops to represent no change in cell state.

To further interpret the predicted cell state transitions, we compared transition probabilities to the change in percent abundance of each cell state and defined a stability metric. We calculate percent abundance for each time point based on the # of cells in a cell state divided by the total number of cells. We defined cell state stability as the percentage of cells not changing state between 8 and 13 months (probability along the diagonal). To isolate condition-specific effects, we further calculated differential stability by subtracting control stability from disease stability.

To provide insight into transcriptome dynamics during AD progression, we interpolated our trajectories based on linear OT paths in cell embedding space to predict expression profiles at time points between 8 and 13 months. Specifically, to retrieve our cell-by-gene expression matrix (*G*) at a certain pseudotime-point, we parameterized all OT-predicted trajectories by a unit-scaled pseudotime value, *t*, between the latent vector representations of our cells, *X_s_* and *X_t_*, and mapped the 100-dimensional cell-by-latent dimension matrix to a 2766-dimensional cell-by-gene matrix (**Equation 3**) via our decoder model. Our pseudo time value (*t*) assumes that linear distance in latent space is aligned with linear timescales (e.g., at *t* = 0.2, our linear interpolation occurs at 20% along the trajectory between our 8 and 13 months cells, assumed to correspond to 9 months in real-time). Thus, although we utilize a pseudotime variable for interpolation, our cell lineages still incorporate true time labels.

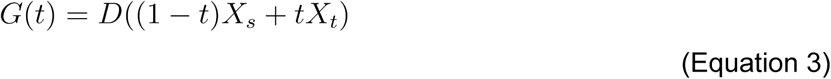

where we calculated the point in latent space, which is a fraction *t* along the OT-mapped trajectory to track gene expression in pseudotime (*t*) between 8 and 13 months. We then reconstructed this point into gene space with our decoder function, *D*.

To identify the relationship between our previously defined spatial DEGs and interpolated samples between 8 and 13 months as predicted by OT, we retrieved the four gene sets, Plaque (UP), Plaque (DOWN), Tau (UP), and Tau (DOWN), and plotted the z-fold change of their median expression in disease and control samples over the interpolated timepoints. We also plot the z-fold change of median DEG expression with respect to each cell state over the interpolated timepoints.

To link increased DEG expression within a cell state to its functional role, we identified the top 50 gene markers for each state within microglia, astrocytes, and oligodendrocytes using the Wilcoxon rank-sum test in Scanpy’s rank_genes_groups method and performed GSEA as described previously.

### Embedded OT for perturbation response modeling

We aimed to predict transcriptome perturbation responses to drug administration via OT[30,103]. Perturbations involved an 80% reduction or increase in target gene expression, simulating knockdown or overexpression, respectively. By conducting perturbations at both 8 and 13 months, we aimed to approximate the effects of sustained treatment over disease progression. We then embedded the perturbed cells into our latent space using our encoder model and calculated the unbalanced OT mapping as described previously. We quantified perturbation-induced cell state changes via net transition matrices, calculated as the perturbed matrix minus the disease matrix, and differential stability, calculated as perturbed stability minus disease stability (**Figure S6**).

Given the spatio-temporal enrichment of the four gene sets (Plaque UP, Plaque DOWN, Tau UP, Tau DOWN), we perturbed potential therapeutic target genes by applying knockdown or overexpression to the top 15 genes in each set. Specifically, we applied knockdown to overexpressed sets (Plaque UP and Tau UP) and overexpression to underexpressed sets (Plaque DOWN and Tau DOWN). We also applied the ChEA3 software[104] for transcription factor (TF) enrichment analysis, identifying key regulatory TFs for each gene set and predicting whether their activation or inhibition aligned with the direction of perturbation. To validate our OT-predicted transition matrices, we cross-referenced each predicted perturbation-induced change in cell state abundance with published experimental evidence, marking matrix entries with supporting citations where increases or decreases had been previously observed.

As a proof-of-concept, we also modeled perturbations based on Donepezil, a common Alzheimer’s treatment that is predicted by DrugBank[105] to inhibit several proteins (**Table S8**)[62,106,107]. Genes coding for these proteins were perturbed to simulate Donepezil treatment.

### Statistics

For computational and statistical analysis, we used Python version 3 and Google Colaboratory[108]. For visualization, we used Matplotlib[87] and Seaborn[93]. The p-values were computed from two-sided tests unless otherwise specified.

## Acknowledgments

We acknowledge general support from National Institutes of Health (NIH) grants AG054012, AG058002, MH109978, AG062377, AG081017, NS129032, AG077227, NS110453, NS115064, AG062335, AG074003, NS127187, AG067151, MH119509, HG008155, DA053631, and DA054584 (to M.K.). This work was supported in part by the Japan Science and Technology Agency PRESTO program, grant number JPMJPR25K7 (to Y.T.). We thank Patricia Purcell, Amy Grayson, the members of the Kellis lab, and Pinar Demetci for their scientific suggestions. The content is solely the responsibility of the authors.

## Data Availability

The analyses presented in this study were based on the single-cell transcriptomics data accessed through the publicly available Broad Institute Single-Cell Portal (https://singlecell.broadinstitute.org/single_cell/) from study SCP1375.

## Code Availability

Analysis scripts in Python will be released upon publication.

## Author Contributions

I.R. conceived the study; Y.T. supervised the study; I.R. designed the study, developed the computational framework, and conducted data analysis with feedback from Y.T.; I.R. wrote the manuscript with feedback from Y.T. and M.K.

## Notes

https://singlecell.broadinstitute.org/single_cell/study/SCP1375/integrative-in-situ-mapping-of-single-cell-transcriptional-states-and-tissue-histopathology-in-an-alzheimer-disease-model

### Competing Interest Statement

Y.T. holds a visiting Associate Professorship at Kyoto University and a visiting researcher position at the University of Tokyo for collaboration; those affiliations have no role in study design, data collection, data analysis, the decision to publish, or the preparation of the manuscript.

