## Supplemental Text & Figures for "sc4D: spatio-temporal single-cell transcriptomics analysis through embedded optimal transport identifies joint glial response to Alzheimer’s disease pathology"

### Supplementary Information

#### Supplementary Figures

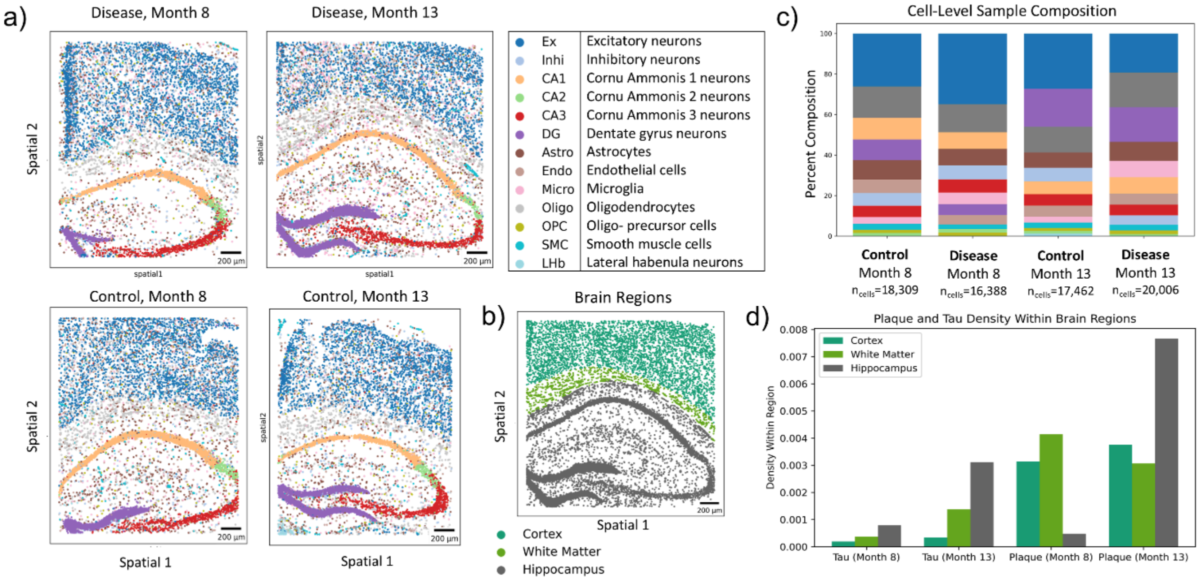


**Supplemental Figure 1. Spatial and longitudinal mapping of sampled brain tissue captures cell type and pathological aggregate localization and progression**. Here, we focus on spatially-resolved distributions of (**a**) cell types in coronal brain slices ($n_{samples}$ = 4) from the hippocampal and cortical regions of control (bottom) and AD+ (top) mice at 8 (left) and 13 months (right) post-induction. 13 pre-labeled major cell types include cortical excitatory neurons (Ex), inhibitory neurons (Inhi), Cornu Ammonis neurons from regions 1, 2, and 3 (CA1, CA2, CA3), dentate gyrus neurons (DG), astrocytes, (Astro), endothelial cells (Endo), microglia (Micro), oligodendrocytes (Oligo), oligodendrocyte precursor cell (OPC), smooth muscle cell (SMC), and lateral habenula neurons (LHb) (**Table S1**). For each mouse brain sample, we report the (**b**) percent composition of each cell type along with the total cell count. In addition, we show the percentage composition of (**c**) pre-labeled cortical, white matter, and hippocampal regions along with the (**d**) spatial distribution of Aβ plaque (top) and PHF+ tau protein (bottom) across our AD+ samples, with bounding boxes indicating regions of high pathological aggregate accumulation. An overlay of spatially resolved cells from (a) is applied. Scale bar, 200 μm.

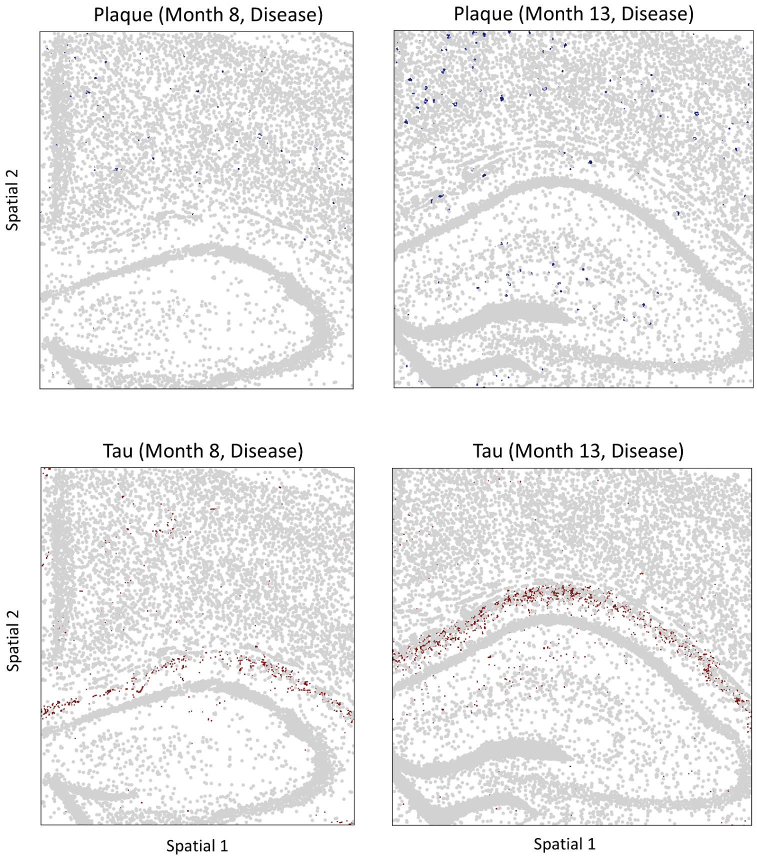


**Supplementary Figure 2. Spatial mapping of plaque and tau across samples**. Here, we focus on spatially-resolved distributions of plaque (blue) and tau (red) aggregates at 8 (left) and 13 (right) months post-induction. We overlay all cells (grey) to provide spatial context for the aggregates.

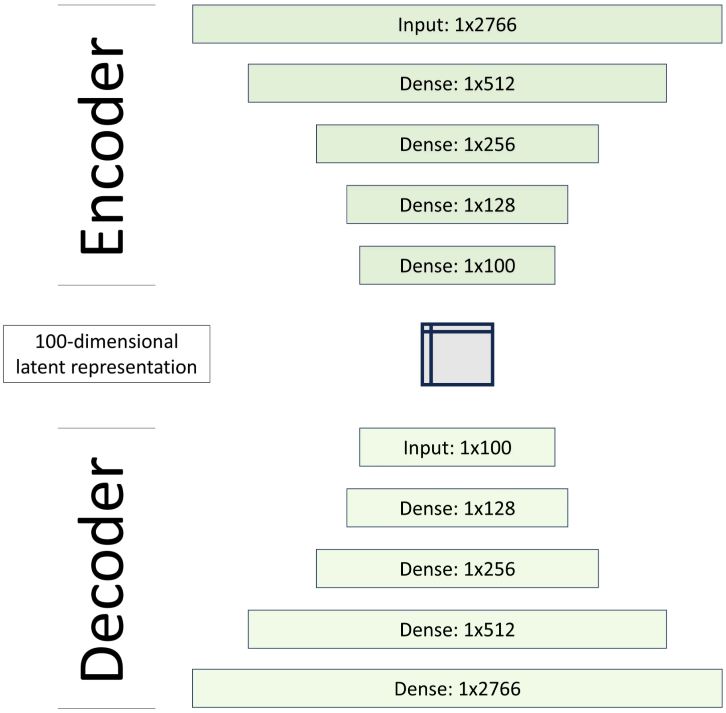


**Supplementary Figure 3. Schematic of autoencoder network architecture.** Encoder and decoder model architecture with one input and four dense layers, annotated by the input shape for each. The encoder embeds single-cell data into a 100-dimensional latent representation, and the decoder reconstructs the gene expression profile from the latent representation.

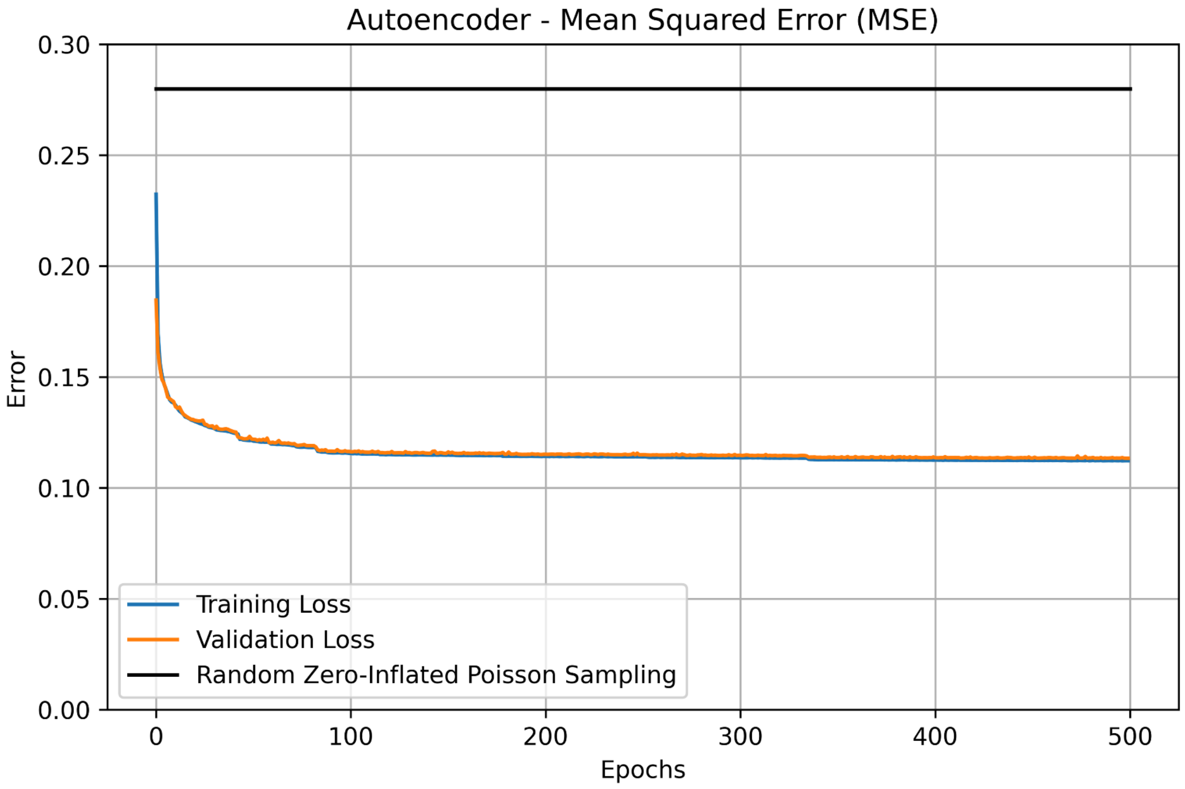


**Supplementary Figure 4. Mean squared error (MSE) training and validation loss across 500 epochs of autoencoder training vs. baseline random sampling model.** The baseline model randomly samples from a zero-inflated Poisson distribution, which closely represents scRNA-seq data distributions[109], yielding a loss of 0.2799. Our autoencoder model achieved a reconstruction training loss of 0.121, a validation loss of 0.122, and a held-out test loss of 0.057.

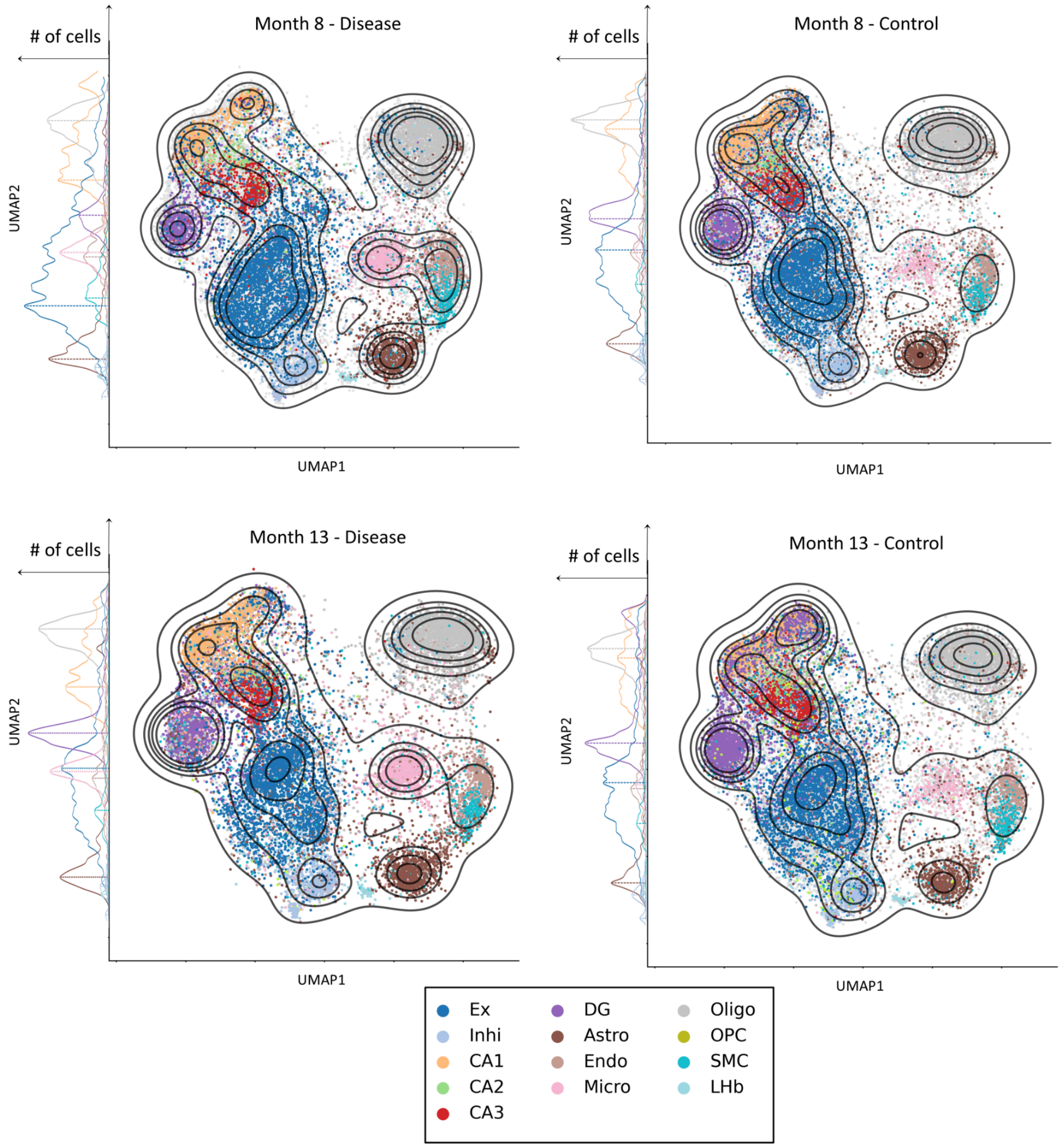


**Supplementary Figure 5. Single-cell UMAP embeddings across all samples.** UMAP embedding of cells from the autoencoder-derived 100-dimensional latent space for our disease (top) and control (bottom) samples between 8 (left) and 13 (right) months post-induction. Kernel density estimation is used to topographically display the distribution of cell observations, which are clustered together and color-coded by cell subtype. We visualize shifts in our cell distributions based on counts of top-level cell types in our latent dimension (UMAP2). A concise version of the supplemental is seen in Figure 5.

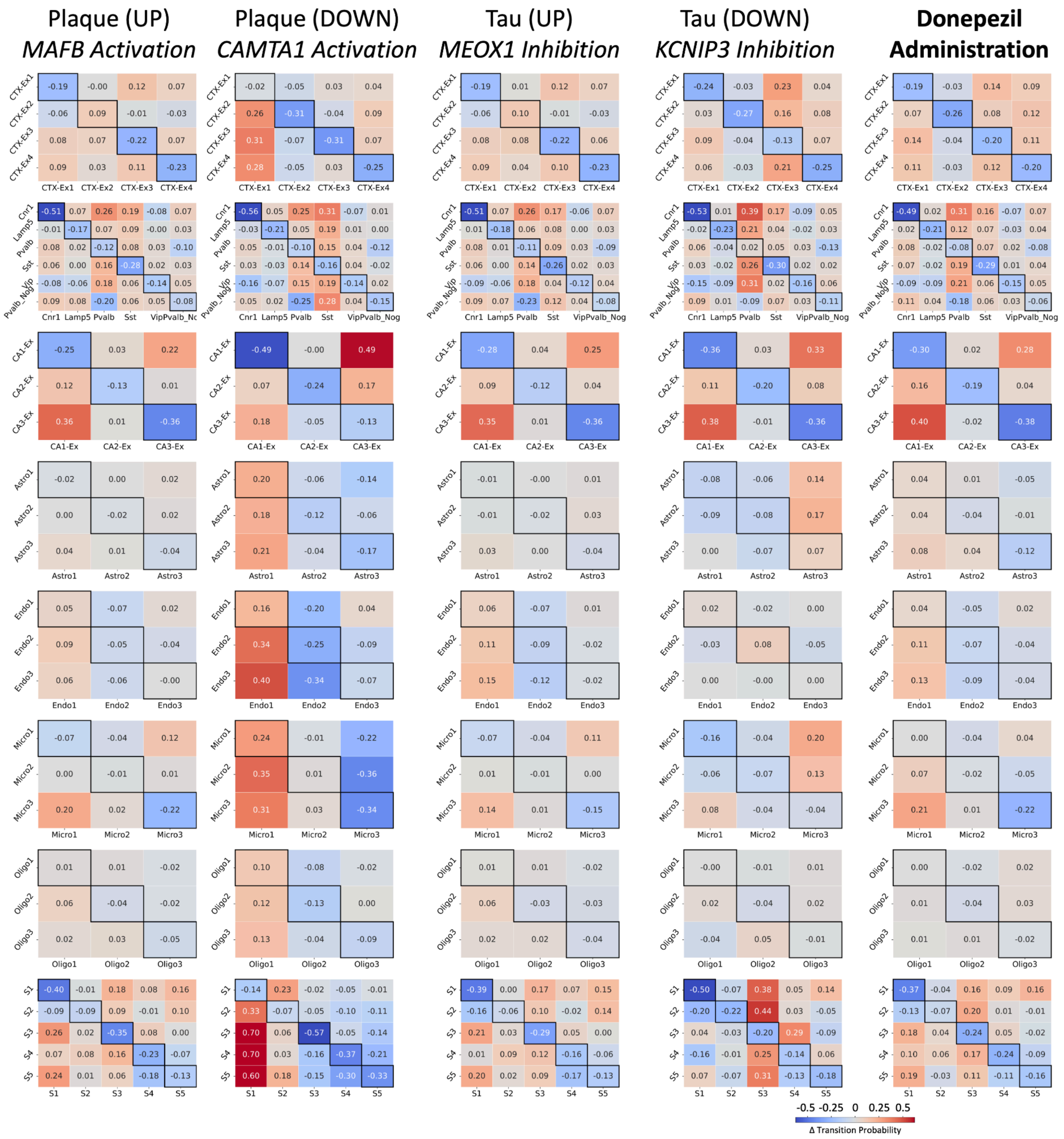


**Supplementary Figure 6. OT-predicted transition matrices across in-silico perturbations.** Five *in-silico* gene perturbations were individually applied to our AD+ single-cell samples before trajectory inference based on DEGs and known pharmacogenetic profiles (**Methods**). Enriched transcription factors regulating each gene set are also noted (**Methods**). Net transition matrices and differential stability of each cell state are shown across all perturbations.

#### Supplementary Tables (see corresponding file)

**Supplementary Table 1. Cell-level composition across samples:** Annotation of cell types and subtypes used throughout the work, in addition to cell counts in control 8-month (a), control 13-month (b), disease 8-month (c), and disease 13-month (d) samples.

**Supplementary Table 2. Cell subtype markers:** Top 10 marker genes for each cell subtype annotation used in this work, as identified by the Wilcoxon rank-sum test in Scanpy's 'rank_genes_groups'.

**Supplementary Table 3. Cell-level spatial composition surrounding pathological aggregates:** Percent abundance of each cell subtype within 10 microns of plaque or tau across all samples.

**Supplementary Table 4. Spatially-enriched DEGs surrounding pathological aggregates:** All spatial DEGs found to be enriched within a 10-micron radius of plaque ($n_{DEGs}$ = 309) or tau ($n_{DEGs}$ = 118) across all samples, with DEGs defined as having an absolute $\log_2$ fold-change > 0.5 and false discovery rate (FDR) < 5% as identified by the Wald test in PyDESeq2.

**Supplementary Table 5. Enriched gene sets found surrounding pathological aggregates:** Enriched (corrected p-value < 0.05) gene sets over- (UP) and under- (DOWN) expressed within a 10-micron radius of plaque and tau from Gene Ontology Biological Process (**Methods**) based on spatial DEGs found previously (**Table S4**).

**Supplementary Table 6. Temporally-enriched DEGs during AD progression:** All temporal DEGs were found to be differentially expressed between our 8 and 13-month samples in disease samples ($n_{DEGs}$ = 646) and control samples ($n_{DEGs}$ = 92) using the Wilcoxon rank-sum test in Scanpy's 'rank_genes_groups'.

**Supplementary Table 7. Temporally-enriched gene sets during AD progression:** Gene sets found to be enriched (corrected p-value < 0.05) during AD progression from 8 to 13 months in disease samples and control samples.

**Supplementary Table 8. Kolmogorov–Smirnov Statistics for Latent Distribution Shifts Across Cell Types:** Quantified changes in 100-dimensional latent space distributions for each major cell type between 8 and 13 months in control and disease samples. For each cell type, Kolmogorov–Smirnov (KS) tests were applied to compare the latent distributions across timepoints. The table reports the maximum KS statistic observed across the 100 dimensions and its corresponding two-tailed p-value.

**Supplementary Table 9. Glial subtype marker genes:** Top 50 marker genes for astrocyte (Astro), microglia (Micro), and oligodendrocyte (Oligo) subtypes. Marker genes are identified using the Wilcoxon rank-sum test in Scanpy's 'rank_genes_groups'.

**Supplementary Table 10. Enriched gene sets for each glial subtype:** Enriched (corrected p-value < 0.05) gene sets for each glial subtype were identified from Gene Ontology Biological Process based on the top 50 marker genes for each subtype.

**Supplementary Table 11. In-silico perturbation gene sets:** Target genes selected for simulated overexpression or knockdown, based on genes over- or underexpressed proximal to plaque or tau. As a proof-of-concept, we model Donepezil administration based on pharmacogenetic profiles and regulatory network analysis from Drugst.One and DrugBank.
